# Sodium selenite alleviates hyperglycemia-aggravated cerebral ischemic injury by mediating apoptosis through the Fas/FasL signaling pathway

**DOI:** 10.1101/2025.08.28.672788

**Authors:** Lan Yang, Feng Ding, Xida Yin, Bowen Zheng, Yue Chang, Jingwen Zhang, Shuai Zhao

**Author notes:** Corresponding author: Email: Yang Lan; Zhao Shuai. These authors are co-author of first author contributed equally to this work (Lan Yang, Feng Ding, Xida Yin). These authors also contributed equally to this work (Bowen Zheng, Yue Chang, Jingwen Zhang).

## Abstract

**Objective:** To investigate the intervention mechanism of sodium selenite in alleviating hyperglycemia-induced cerebral ischemia/reperfusion (I/R) injury through the Fas/FasL/NF-κB/PUMA pathway.

**Methods:** The SD rat model of hyperglycemia-induced cerebral I/R injury and the HT22 cell model of high-glucose oxygen deprivation/reoxygenation (OD/R) were constructed to systematically evaluate the neuroprotective effect of sodium selenite and its regulatory effect on the Fas/FasL/NF-κB/PUMA pathway.

**Results:** The infarct volume, neurological dysfunction, metabolic level, and apoptosis level in the HG group were significantly higher than those in the NG group. Sodium selenite improved the infarct volume, neurological dysfunction, metabolic level, and apoptosis level in the HG group, and regulated the Fas/FasL/PUMA apoptotic pathway to alleviate injury. In vitro experiments showed that sodium selenite significantly improved morphological damage caused by high-glucose OD/R, increased cell viability, and reduced the expression of apoptotic factors. The application of Fas and FasL inhibitors and knockdown of PUMA expression indicated that sodium selenite alleviated cell apoptosis by inhibiting the activation of the Fas/FasL/PUMA pathway.

**Conclusion:** Sodium selenite alleviates hyperglycemia-exacerbated cerebral I/R injury by regulating the Fas/FasL/NF-κB/PUMA signaling pathway.

## 1. Introduction

Stroke remains the second leading cause of death globally, causing approximately 5.5 million deaths each year **Error! Reference source not found.**. It is also the primary cause of long-term disability worldwide. The latest epidemiological surveillance shows that the incidence of stroke is rising at an alarming rate, and it is predicted that 25% of the global population will experience at least one stroke in their life time [1]. Pathologically, stroke can be divided into two major subtypes: ischemic and hemorrhagic. Studies have confirmed that ischemic stroke accounts for 70-80% of all stroke cases and is the most common type of stroke globally **Error! Reference source not found.**[2]. When ischemic stroke occurs, multiple signaling pathways within brain cells are activated. Insufficient blood supply to the brain and interruption of blood flow lead to a lack of energy in brain cells. The energy-dependent ion pump system stops working, disrupting the ion balance inside and outside the cells. A large amount of excitatory neurotransmitters such as glutamate are released, which are toxic to neurons, resulting in neuronal death [3]-[5].

Diabetes or hyperglycemia has been clearly identified as an independent risk factor for ischemic stroke **Error! Reference source not found.**. Some studies have shown that the severity of stroke is usually related to the degree of hyperglycemia. Neuropathological examinations have found that diabetes causes specific microvascular lesions such as hyperplasia and aggravated hyalinization of cerebral microvascular endothelial cells [7] and accelerated intracranial atherosclerosis [8][10]. These lead to faster ischemia, larger infarct volume, more severe neurological deficits, delayed functional recovery, and increased mortality in diabetic stroke patients **Error! Reference source not found.**. During acute ischemic stroke, hyperglycemia significantly exacerbates the pathophysiological process and clinical consequences of acute ischemic stroke through multiple mechanisms, including energy metabolism disorder, increased oxidative stress, increased vulnerability of the penumbra, damage to the neurovascular unit, and activation of the inflammatory response **Error! Reference source not found. Error! Reference source not found.**.

The Fas/FasL signaling axis is a key apoptotic cascade pathway. Fas is a death receptor that transmits an apoptosis-inducing signal when activated by its ligand FasL **Error! Reference source not found.**. The Fas signaling pathway has been considered a key inducer of apoptotic signals during acute cerebral ischemia [14]. The incidence of ischemic stroke is highly correlated with the expression of Fas, Fas-associated death domain protein (FADD), and Caspase-8 [16]. Nuclear factor-κB (NF-κB), a key factor downstream of Fas/FasL, is involved in the occurrence and regulation of downstream apoptosis. NF-κB is a ubiquitous inducible transcription factor responsible for mediating the expression of a large number of genes involved in differentiation, apoptosis, and proliferation **Error! Reference source not found.**. PUMA, a cell apoptosis regulator and a downstream factor of NF-κB, is one of the most effective mediators of cell apoptosis [17]. In hepatocytes, the lack of PUMA can block the p53 activation and apoptotic response to DNA damage induced by hypoxia [18]. PUMA can induce mitochondrial dysfunction and Caspase activation through other members of the Bcl-2 family, including Bcl-2, Mcl-1, and Bcl-xL [19]. Studies have shown that the Fas/FasL complex can induce the expression of the pro-apoptotic protein PUMA by regulating the autophagy process and activating the NF-κB signaling pathway, ultimately leading to programmed death of hepatocytes [20]. However, the relationship between the Fas/FasL/NF-κB/PUMA signaling pathway and hyperglycemia-aggravated cerebral I/R injury remains unclear.

Sodium Selenite (SE) is an important selenium compound and a key trace element for maintaining the functions of human and animal cells **Error! Reference source not found.**. Its mechanism of action is involved in various physiological regulations. Studies have shown that SE significantly increases the expression of SE-dependent enzymes such as glutathione peroxidase [22], antagonizes the central toxicity of heavy metals such as methylmercury and lead, and neurotoxins [23], and plays an important role in immune regulation, mammalian development, and male reproductive function [24]. In cell studies, SE enhances the antioxidant system by activating superoxide dismutase (SOD) and glutathione peroxidase (GPx) [25], reduces glutamate toxicity, metal-induced oxidative damage, and inflammatory cytokine attacks [26][28], and inhibits cell apoptosis by regulating the Bcl-2/Bax protein ratio [29] [31]. SE regulates the MAPK, PI3K-Akt, and NF-κB signaling pathways [32]**Error! Reference source not found.**, activates mitochondrial regeneration signals to maintain the integrity of mitochondrial function [34], and plays a protective role in the cardiovascular system by inhibiting arteriosclerosis and metal toxicity **Error! Reference source not found.**. Research has shown that pretreatment with SE can reduce the infarct area after cerebral ischemia [34], alleviate hyperglycemia-aggravated reperfusion injury **Error! Reference source not found.**, and effectively reduce mitochondrial oxidative stress in Parkinson’s disease and hippocampal ischemia models [35] [37]. However, its neuroprotective mechanism in the Fas/FasL/NF-κB/PUMA signaling pathway remains to be elucidated.

Based on the above, we used a hyperglycemic cerebral I/R animal model and a high-glucose OD/R cell model to study whether SE protects against hyperglycemia-aggravated cerebral I/R injury through an anti-apoptotic mechanism and to explore whether its regulatory mechanism is related to the Fas/FasL/NF-κB/PUMA signaling pathway.

## 2. Materials and methods

### 2.1 Animals and grouping

This study was approved by the Animal Ethics Committee of Ningxia Medical University (Approval No.: IACUC-NYLAC-2023-038). SD rats aged 8 weeks, weighing between 210 g and 230 g, were selected and housed in a specific-pathogen-free environment (air humidity 55% ± 5%, 12h light/12h dark cycle, room temperature 25°C, with free access to water and food). After transient middle cerebral artery occlusion (MCAO) surgery, the rats were finally divided into four groups: normoglycemic sham-operated group (Sham; n = 10), normoglycemic cerebral ischemia/reperfusion group (NG+I/R; n = 10), hyperglycemic cerebral ischemia/reperfusion group (HG+I/R; n = 10), and SE-intervened hyperglycemic cerebral ischemia/reperfusion group (HG+I/R+SE; n = 10).

### 2.2 Animal models and treatment methods

#### Establishment of the Hyperglycemic model

1% streptozotocin (STZ, S8050, Solarbio, China) was dissolved in phosphate-buffered saline (PBS) buffer (C3580-P050K, Biological Industries, Israel) for injection. Rats in the HG group and the HG+SE group were intraperitoneally injected with STZ (60 mg/kg). After 72 h, rats with fasting blood glucose (FBG) levels higher than 16.7 mM were defined as hyperglycemic. Rats in the NG group were treated with PBS buffer according to their body weight.

#### Establishment of the MCAO model

After 4 weeks of SE treatment, the MCAO model was established using a modified thread-embolization method. Rats were deeply anesthetized with isoflurane (3% for induction, 2% during surgery). A 3 cm surgical incision was made in the neck to isolate and expose the right common carotid artery, internal carotid artery, and external carotid artery. In the experimental groups, the proximal end of the common carotid artery and the origin of the external carotid artery were double-ligated. Then, a 1 mm oblique incision was made at the distal end of the common carotid artery, and a thread was inserted and advanced to 18-20 mm from the bifurcation of the common carotid artery. Insertion was stopped when resistance was felt and the marked point reached the incision, and the thread was double-ligated and fixed. The thread was removed after 30 min, and the skin was sutured layer by layer. After surgery, rats were placed on an electric blanket to maintain body temperature and returned to their cages after waking up. In the Sham group, only the skin was incised and the blood vessels were isolated. The blood vessels were not cut, and no thread was inserted. Other operations were the same as those in the surgical group.

#### Sodium Selenite intervention

Sodium selenite (#214485, SIGMA-ALDRICH, USA) was dissolved in PBS to prepare a 0.04 mg/mL solution. After the hyperglycemic model was established, rats in the HG+SE group were intraperitoneally injected with SE (0.4 mg/kg/d) for 4 weeks.

### 2.3 2,3,5-Triphenyltetrazolium Chloride (TTC) Staining

In this study, two methods were used to analyze the brain damage caused by reperfusion after MCAO treatment. TTC staining was used to evaluate the volume of cerebral infarction. The modified neurological severity score (mNSS) method was used to evaluate neurological deficits. Briefly, the whole brains of rats were cut into slices approximately 4 mm thick along the coronal plane and then stained with a TTC solution (1%) at 37°C for 30 min. The infarcted area was white, and the non-infarcted area was red. The infarct volume was calculated by measuring the area of the infarcted region in each slice after TTC staining, multiplying it by the slice thickness, dividing the infarct volume by the total volume of the brain slices, and then multiplying by 100% to obtain the percentage of infarct volume.

### 2.4 Neurological deficit assessment

For short-term neurological deficit assessment, the mNSS test (maximum score of 18 points), including motor, sensory, balance, reflex, and muscle tone, was performed by two researchers who were blind to the experiment. A damage score between 13 and 18 points indicated severe damage, 7 to 12 points indicated moderate damage, and 1 to 6 points indicated mild damage.

### 2.5 Treatment of Paraffin-Embedded specimens

Anesthetized rats were decapitated to obtain their brains. For paraffin-embedded specimens, brain slices were soaked in 4% formaldehyde solution for more than 72h, placed in paraffin-embedding cassettes, and then the cassettes were placed in an automatic tissue dehydrator for dehydration.

### 2.6 Hematoxylin-Eosin (HE) staining and Nissl staining

Similarly, paraffin-embedded brain tissues fixed with 4% formaldehyde were prepared into 4 μm thick slices. After standard dewaxing and gradient ethanol hydration of the paraffin slices, HE staining (G1129, Solarbio, China) and Nissl staining (G1430, Servicebio, China) were performed.

#### HE staining

First, for dewaxing and dehydration, after baking the slices at 65°C for 1 h, the slices were soaked in xylene I (20 min), xylene II (15 min), absolute ethanol I (10 min), absolute ethanol II (10 min), 95% alcohol (5 min), 85% alcohol (5 min), 75% alcohol (5 min) in sequence, and then rinsed with distilled water to remove impurities from the slices. Then, hematoxylin staining was carried out. The dewaxed and dehydrated slices were stained in hematoxylin staining solution for 10 min and rinsed with running distilled water until clear, making the cell nuclei blue. Subsequently, differentiation and bluing back were performed. The slices after hematoxylin staining and washing were differentiated in a hydrochloric acid-alcohol mixture for 3 s and immediately rinsed with distilled water, changing the cell nuclei from dark blue to light blue. Then, eosin staining was carried out. The slices after differentiation, bluing back, and washing until clear were stained in eosin staining solution for 45 s and rinsed with distilled water until clear, making the cytoplasm and extracellular matrix red or pink and the cell nuclei light blue. Finally, for transparency and mounting, the slices after eosin staining and washing were soaked in 75%, 85%, 95% alcohol (5 min each), absolute ethanol I (10 min), absolute ethanol II (10 min), xylene I (15 min), xylene II (15 min) in sequence, mounted with neutral gum, and observed and imaged under a microscope after drying. HE staining was used to evaluate the proportion of apoptotic nerve cells.

#### Nissl staining

Dewaxing and dehydration are the same as in HE staining. Then, place the tissue sections into a staining jar filled with pre-prepared Nissl working solution. Next, put the staining jar in an oven at 60°C and let the samples incubate in a stable temperature environment for 1 h. After incubation, take the staining jar out of the oven and rinse the samples twice with distilled water to remove the unbound excess dye. Then, perform differentiation. Prepare an alcohol-hydrochloric acid mixture and quickly place the stained and washed samples into it for differentiation. During the differentiation process, observe the staining of the samples under a microscope in real-time. Adjust the differentiation time to ensure that the morphology and structure of Nissl bodies can be clearly observed, making the Nissl bodies show obvious and clear morphological characteristics under the microscope. Finally, perform clearing and mounting, which is the same as in HE staining. Evaluate the number of Nissl bodies through Nissl staining.

### 2.7 Transmission electron microscopy detection

Collect fresh brain tissue from the junction between the right infarct area and normal tissue. Cut the fresh brain tissue into a volume of 6 mm³. Fix it with 3% glutaraldehyde fixative and 0.1% osmium acid. Then, wash, dehydrate, embed the specimens in epoxy resin, make ultrathin sections, and stain them with 3% uranyl acetate and lead citrate. Finally, observe and take pictures using a transmission electron microscope (HITACHI HT7800, Japan).

### 2.8 Immunohistochemical staining

We perform dewaxing, antigen retrieval, blocking, primary antibody incubation, secondary antibody incubation, color development reaction, counter-staining, clearing, and mounting on paraffin sections of rat brain tissue in sequence. First, soak the sections in xylene for 20 min (twice, 10 min each time), then soak them in absolute ethanol, 95% ethanol, 85% ethanol, and 75% ethanol in sequence (5 min each), and finally rinse them with distilled water to remove impurities. Then, place the sections in 0.01 mmol/L citrate retrieval solution and perform high-pressure retrieval in a pressure cooker for 10 min. After cooling, rinse them with PBS three times (5 min each time). Circle the tissue on the sections with an immunohistochemical pen, add 0.3% hydrogen peroxide solution, and incubate at room temperature for 15 min. After rinsing with PBS three times, add goat serum working solution and incubate at room temperature for 30 min. Then, add the diluted primary antibody (1:200-1:500) and incubate at 4°C overnight. After rewarming at room temperature for 1 h, rinse with PBS three times, add the secondary antibody (rabbit-anti or mouse-anti) and incubate at room temperature for 30 min, and then rinse with PBS three times. Add DAB color-developing working solution, observe the color developing situation under a microscope, and stop the reaction in time. Rinse with PBS three times. Add hematoxylin solution for counter-staining for 5 min, differentiate with hydrochloric acid-alcohol for 3 s, and wash with water to turn blue. Finally, soak the sections in 75% ethanol, 85% ethanol, 95% ethanol, absolute ethanol, and xylene in sequence (5 min each), mount them with neutral gum, and observe under a microscope after drying.

### 2.9 Cell culture and drug preparation

The HT22 cell line (mouse hippocampal neurons) is from Hysigen Co., Ltd. (Hysigen, China, No. #TCM-C821). The 3MA culture medium is from Absin (abs810575, Absin, China). Culture HT22 cells in DMEM/F12 medium (#C3130-0500, VivaCell, China) containing 10% fetal bovine serum (A5669701, Thermo Fisher, China) and 1% penicillin/streptomycin (#03-031-5B, Biological Industries, Israel). Culture the cells in an incubator at 37°C with 5% CO₂. The DMEM/F12 medium originally contains 17.5 mM glucose and is regarded as a medium with normal glucose concentration. According to previous studies [38], a medium with high-glucose concentration is defined as 50 mM. Therefore, we select a glucose concentration of 50 mmol/L to prepare a high-glucose medium. Dissolve 1.35 g of glucose powder in 5 mL of complete HT22 cell culture medium to prepare a high-glucose stock solution. Prepare a SE stock solution with a concentration of 100 nmol/L in the same way. Weigh the SE powder, add it to the complete culture medium, and stir it continuously with a magnetic stirrer until it is completely dissolved to form a clear and transparent stock solution.

### 2.10 High-glucose OD/R model and grouping

Inoculate the cells into the culture medium according to the experimental groups and perform routine culture for 24 h. When the cell density reaches 70%, aspirate the original culture medium. Rinse the cells twice with PBS, and then replace it with a 50 mmol/L high-glucose medium. Immediately put the culture dish, an anaerobic pack (#A-7/D-7, Mitsubishi Chemical, Japan), and an oxygen indicator (#C-22/D-66, Mitsubishi Chemical, Japan) into a 2.5-liter sealed box. Then, put the sealed box into an incubator at 37°C. After 1 h, when the color of the oxygen indicator changes from dark purple to light pink, an anaerobic environment with an oxygen concentration of less than 0.1% and a carbon dioxide concentration of 5% is achieved in the sealed box. Record this time as the start time of the OD experiment. After 4 h, open the sealed box and transfer the cells to a normal incubator (5% CO₂, 95% air) for 24 h reoxygenation culture to construct a cell model of hyperglycemia combined with cerebral I/R injury.

In in vitro experiments, divide the HT22 cells into a CON group, a normal-glucose oxygen deprivation/reoxygenation group (NG+OD/R), a high-glucose oxygen deprivation/reoxygenation group (HG+OD/R), a high-glucose oxygen deprivation/reoxygenation+SE group (HG+OD/R+SE), and a high-glucose oxygen deprivation/reoxygenation+SE+3MA autophagy inhibitor group (HG+OD/R+SE+3MA).

Divide the HT22-PUMA gene-knocked-down cells and their control cells into a CON group, a high-glucose oxygen deprivation/reoxygenation group (HG+OD/R), and a high-glucose oxygen deprivation/reoxygenation+SE group (HG+OD/R+SE).

### 2.11 CCK-8 assay

According to the manufacturer’s protocol, the CCK-8 kit (CK04, DOJINDO, Japan) was used to evaluate cell viability. Briefly, cells in the logarithmic growth phase were seeded and cultured in 96-well plates (100 μL, 2×10^3^ cells per well). After the OD/R experiment, 10 μL of CCK-8 reagent was added to each well, and the cells were incubated again at 37°C for 2 h. The absorbance value of each well was measured at 532 nm using a microplate reader.

### 2.12 Western blot

At two time points of 24 h and 72 h after I/R in rats, the rats were anesthetized and their brains were removed. 100 mg of brain tissue from the ischemic penumbra area was excised for subsequent experiments. After the cells were subjected to OD/R intervention, the culture medium was aspirated, and the cells were washed with PBS. The cells were scraped off with a cell scraper and transferred to a centrifuge tube. After centrifugation at 2500 rpm for 5 min, the supernatant was aspirated, and the lysis buffer was prepared and added according to the instructions. Total protein was extracted from tissues or cells using a cell lysis buffer kit (#P0013, Beyotime, China). Proteins of different molecular weights were separated by SDS-polyacrylamide gel electrophoresis. Then the proteins were transferred to a polyvinylidene fluoride (PVDF) membrane (Millipore, USA). First, the membrane was immersed in 5% non-fat milk powder or 3% bovine serum albumin and incubated for 2 h. Next, the membrane was washed thoroughly with TBST three times and then immersed in the primary antibody solution and incubated overnight at 4°C. The primary antibodies were as follows: PUMA Rabbit mAb (#24633, CST, USA), Cytochrome c Rabbit mAb (#4280, CST, USA), Cleaved Caspase-3 Rabbit mAb (#9661, CST, USA), Bax Rabbit mAb (#41162, CST, USA), Bcl-2 Mouse mAb (#sc-7382, Santa, USA), Rabbit anti-Fas Ligand Polyclonal Antibody (abs131431, Absin, China), Rabbit anti-FAS Polyclonal Antibody (abs158773, Absin, China), NF-kappa B p65 Rabbit mAb (#8242), Phospho-NF-kappa B p65 Rabbit mAb (#3033), βeta-Actin (Loading Control) Rabbit pAb (#bs-0061R, Biosynthesis, China). The next day, after the membrane was washed thoroughly with TBST, it was immersed in goat anti-mouse IgG (H+L) secondary antibody (#abs20040, Absin, China) or goat anti-rabbit IgG (H+L) secondary antibody (#S0002, Affinity, USA) and incubated at room temperature for 2 h. Finally, the blot bands were visualized using ECL reagent.

### 2.13 Quantitative and statistical analysis

The image information was quantitatively analyzed by ImageJ software (Fiji is ImageJ 2.1.0/1.53q/Java 1.8.0_172). The statistical charts of the experimental data were drawn by GraphPad Prism 8 software. The graphical abstract was drawn by Figdraw. All the data obtained in the experiment were statistically processed by IBM SPSS Statistics 20 software. All measured data were expressed as mean ± standard deviation (SD). The statistical differences between the means of multiple independent samples were analyzed by one-way analysis of variance (One-Way ANOVA), followed by pairwise comparison between groups using the LSD method. P < 0.05 was considered statistically significant.

## 3. Results

### 3.1 Successful establishment of the hyperglycemia model

In Fig 1a, we presented the body characteristics of rats under different treatments. In NG group, the rats appeared plump, with smooth and healthy-colored fur, indicating good nutritional status. In contrast, the rats in HG group treated with STZ were thinner, with rough and yellowish fur, reflecting poor nutritional status. In HG+SE group, the body posture, fur color, and overall nutritional status of the rats showed a positive improvement trend compared with the simple HG group. Fig 1b shows the HE staining of rat pancreas. In NG group, the pancreatic structure of the rats was intact and the pancreatic morphology was normal. However, in HG group and HG+SE group, the pancreas was significantly damaged, the islets were irregular in shape, and the invasion and hyperplasia of exocrine acinar cells could be seen. Fig 1c shows the immunofluorescence results of rat islet A cells. Compared with the NG group, the glucagon expression in the rats of the HG group and the HG+SE group was decreased, and the islets were significantly smaller and irregular in shape.

**Fig 1.**
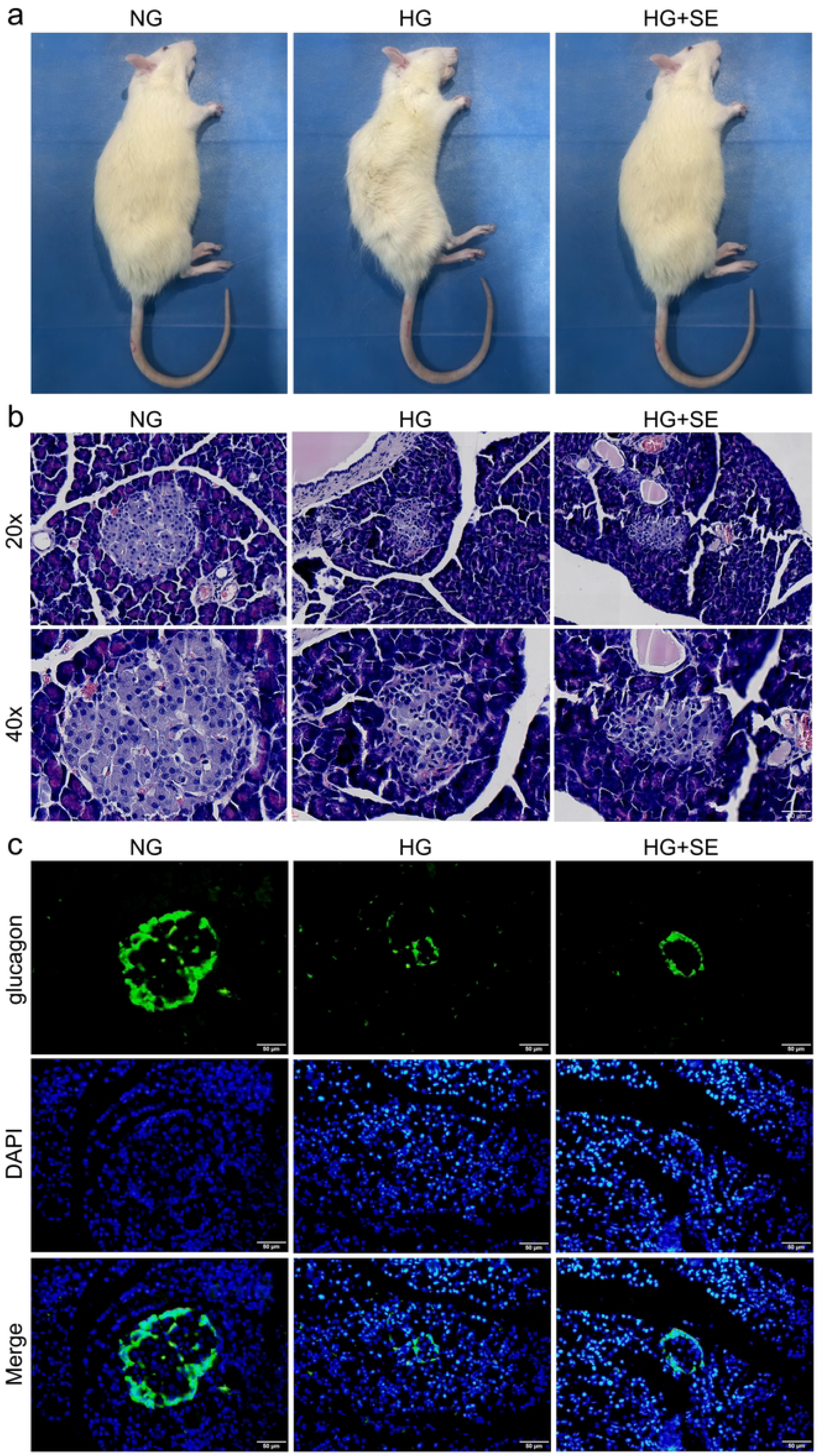
The hyperglycemia model was successfully constructed. **a** Appearance characteristics of the rats. **b** HE staining results of the rat pancreas. The first row shows the pancreatic tissue observed under a 20× microscope, and the second row shows the pancreatic tissue observed under a 40× microscope (scale bar = 50μm). **c** The immunofluorescence results of glucagon in the rat islets (scale bar = 50μm).

### 3.2 SE alleviates the metabolic level, neurological dysfunction, and infarct volume in rats with cerebral I/R aggravated by hyperglycemia

In Fig 2, the representative TTC staining images (Fig 2a and 2b) show that hyperglycemia significantly increased the cerebral infarct volume (white area) after I/R injury (P<0.05), while SE treatment significantly reduced the infarct volume (P<0.05). The neurological deficit scores after cerebral I/R injury in Fig 2c indicate that at the two time points of 24 h and 72 h, the scores of the rats in the HG group and the HG+SE group were higher than those of the rats in the NG group (P<0.05), and the scores of the rats in the HG+SE group were significantly lower than those of the rats in the HG group (P<0.05). Hyperglycemia significantly aggravated the neurological damage in the body, while SE could effectively alleviate this damage. The body weight, blood glucose level, and water, urine, and food intake of the rats were continuously monitored for more than four weeks. As shown in Fig 2 d-f, the metabolic rate of the diabetic rats was significantly higher than that of the other groups (P<0.05), the body weight gain was extremely slow, and the average fasting blood glucose was always higher than 16.7 mM.

**Fig 2.**
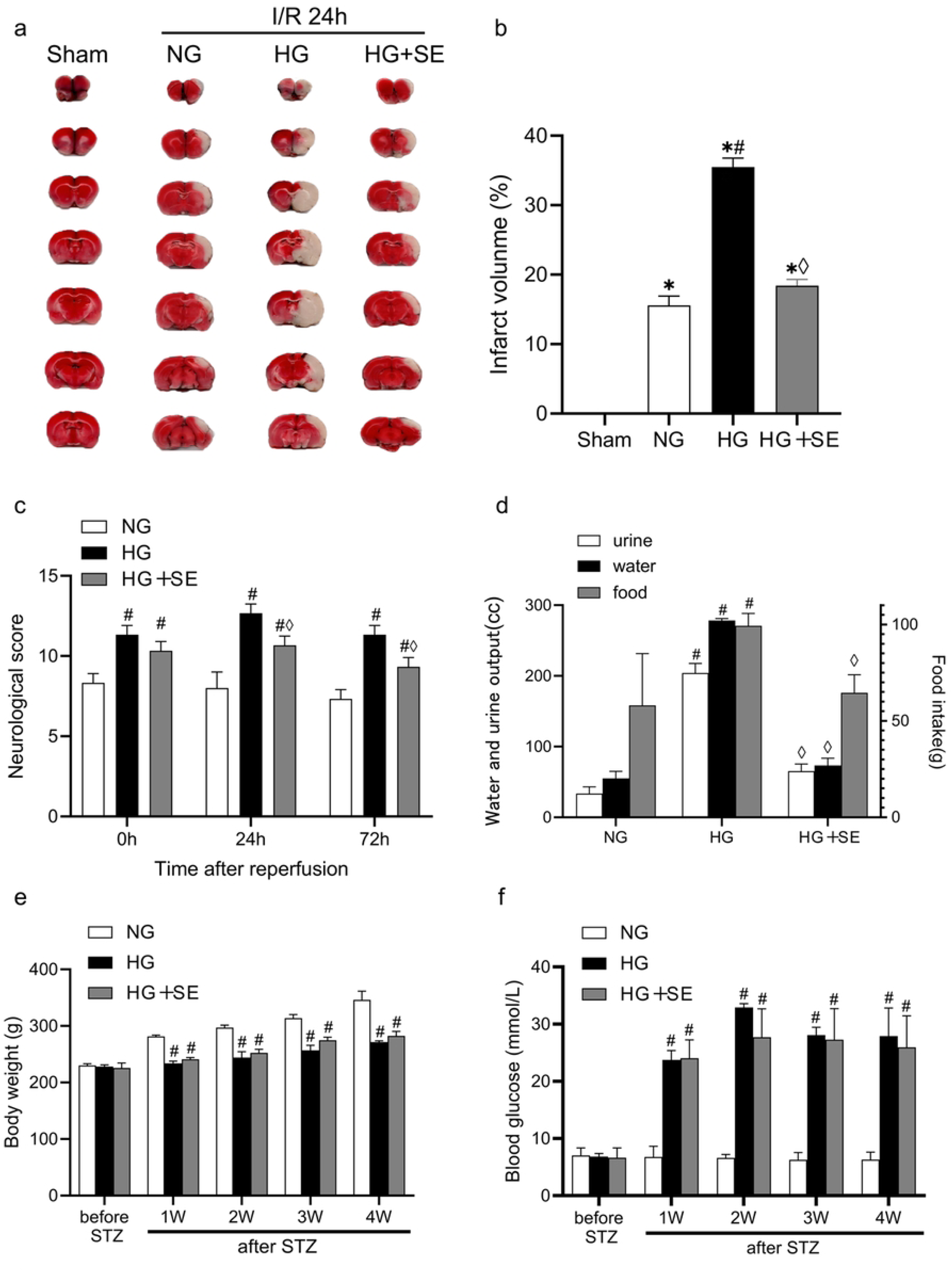
SE alleviates the infarct volume, neurological dysfunction, and metabolic level aggravated by hyperglycemia in rats with cerebral I/R. **a** TTC staining. **b** The statistical results of infarct volume by TTC staining (n = 3). **c** Neurological deficit scores in rats (n = 20). **d** Metabolic rate measurement in rats at 24 h. **e** Body weight changes in rats. **f** Blood glucose changes in rats. *P < 0.05 compared with the Sham group; ^#^P < 0.05 compared with the NG+I/R group; ^◊^P < 0.05 compared with the HG+I/R group.

### 3.3 SE improves neuronal damage in rats with cerebral I/R aggravated by hyperglycemia

We performed HE staining on the rat brain tissue. The results (Fig 3a-c) showed that compared with the rats in the Sham group, different degrees of neuronal damage occurred in the brain tissue of the rats in the NG group, the HG group, and the HG+SE group after MCAO modeling. At 24 h after I/R, compared with the NG group, HG group showed more severe nuclear pyknosis and neuronal edema; at 72 h after I/R, the damage was alleviated compared with that at 24 h. Compared with HG group, the morphological changes mentioned above were alleviated in the HG+SE group at both 24 h and 72 h after I/R.

**Fig 3.**
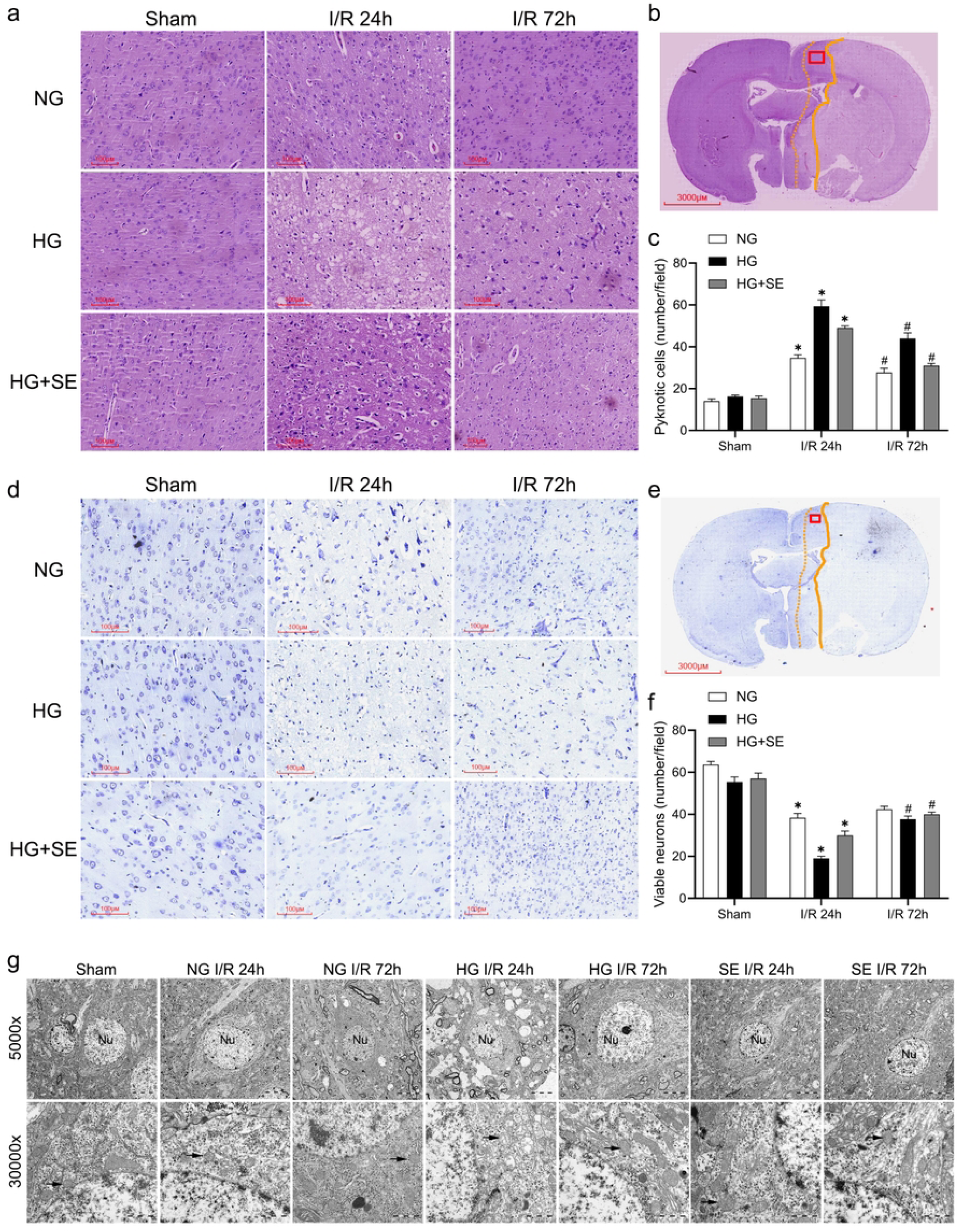
SE improves neuronal damage in rats with cerebral I/R exacerbated by hyperglycemia. a and b HE staining of brain tissue. **a** Representative HE-stained section of the infarct border zone (ischemic penumbra) from the area marked by the red box in b (scale bar = 100 μm). **b** Panoramic scan of HE-stained cerebral section (scale bar = 3 mm). **c** Quantitative analysis of necrotic neurons in HE-stained sections (n = 3). **d** Representative Nissl-stained section of the infarct border zone (ischemic penumbra) from the area marked by the red box in e (scale bar = 100 μm). **e** Panoramic scan of Nissl-stained cerebral section (scale bar = 3 mm). **f** Quantitative analysis of neurons with surviving Nissl bodies (n = 3). **g** Transmission electron microscopy of ischemic penumbra: First row micrographs at ×5000 magnification (scale bar = 5 μm). Second row micrographs at ×30000 magnification (scale bar = 1 μm). *P < 0.05, compared with the Sham group; ^#^P < 0.05, compared with the NG+I/R group.

Fig 3d-f shows the results of Nissl staining of the rat brain. The neuronal morphology in the Sham group remained normal, and the Nissl bodies were clearly distinguishable. At 24 h after I/R, compared with the NG group, the neurons in the HG group showed nuclear pyknosis, and the Nissl bodies became blurred or even completely disappeared. This damage was alleviated at 72 h after I/R. Compared with the HG group, the morphological changes such as nuclear pyknosis and blurring or disappearance of Nissl bodies were significantly alleviated in the HG+SE group at both 24 h and 72 h after I/R.

Subsequently, electron-microscopic examination was performed on the ischemic penumbra of the rat brain (Fig 3g). In the Sham group, the cell-membrane structure in the ischemic penumbra of the brain was intact, and the organelles such as mitochondria, endoplasmic reticulum, and Golgi apparatus were intact, and the nuclear-membrane structure was intact. Compared with the Sham group, varying degrees of nuclear-membrane wrinkling, increased heterochromatin in the nucleus, decreased number of mitochondria, mitochondrial swelling, disappearance of cristae, and increased density of mitochondrial matrix occurred in the NG group, the HG group, and the HG+SE group; the above-mentioned conditions were the most severe in the HG group, with increased heterochromatin in the nucleus and severe vacuolization in mitochondria and cytoplasm. At 72 h after I/R, the damage was alleviated compared with that at 24 h. Compared with the HG group, the morphological changes mentioned above were alleviated in the HG+SE group at both 24 h and 72 h after I/R.

### 3.4 SE alleviates cell apoptosis in cerebral I/R injury exacerbated by hyperglycemia

To investigate the protective effect of selenium against cerebral I/R injury exacerbated by hyperglycemia in vivo, we detected the expression levels of key factors in the pathway (Fas, FasL, PUMA, NF-κB and its phosphorylation) and apoptosis-related factors in the penumbra of the rat brain by Western Blot (Fig 4). As shown in Fig 4a, b/c/e/f, compared with the Sham group, the expression levels of Fas, FasL, PUMA, NF-κB and its phosphorylation in the NG group, the HG group, and the HG+SE group showed an increasing trend. Compared with the NG group, the expression levels of Fas and p-NF-κB in the HG group were significantly increased at 24 h after I/R (P<0.05). At 72 h after I/R, the expression levels of Fas and FasL in the HG group were significantly increased (P<0.05). Compared with the HG group, the expression levels of Fas, FasL, PUMA, NF-κB and its phosphorylation in the HG+SE group were significantly decreased at both 24 h and 72 h after I/R (P<0.05).

**Fig 4.**
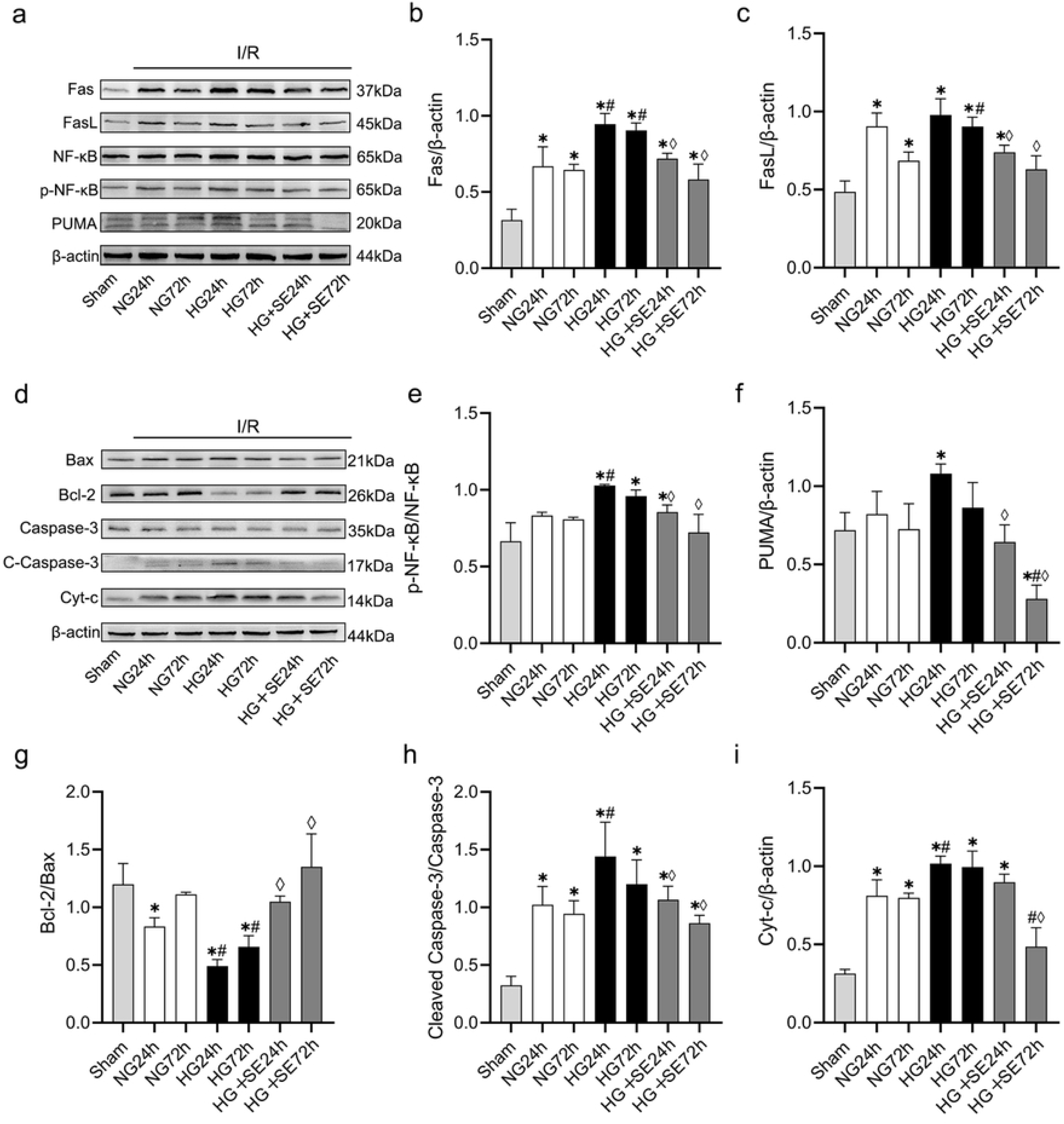
SE alleviates hyperglycemia-exacerbated apoptosis in rats with cerebral I/R injury. **a** Representative Western Blot results of Fas/FasL/NF-κB pathway proteins. **b** Quantitative analysis of Fas protein expression (relative to β-actin) from a (n=3). **c** Quantitative analysis of FasL protein expression (relative to β-actin) from a (n=3). **e** Quantitative analysis of p-NF-κB activation level (p-NF-κB/NF-κB ratio) from a (n=3). **f** Quantitative analysis of PUMA protein expression (relative to β-actin) from a (n=3). **d** Representative Western Blot results of apoptosis-related proteins in rat brain tissue. **g** Quantitative analysis of Bcl-2/Bax expression from d (n=3). **h** Quantitative analysis of Cleaved Caspase-3/Caspase-3 expression from d (n=3). **i** Quantitative analysis of Cyt-c protein expression (relative to β-actin) from d (n=3). *P < 0.05, compared with the Sham group; ^#^P < 0.05, compared with the NG+I/R group; ^◊^P < 0.05, compared with the HG+I/R group.

The expression levels of apoptosis-related factors Bax, Bcl-2, Cleaved Caspase-3, and Cyt-c in the penumbra of the rat brain are shown in Fig 4d. As shown in Fig 4g/h/i, compared with the Sham group, the expression levels of Cleaved Caspase-3 and Cyt-c in the NG group, HG group, and HG+SE group were significantly increased, while the Bcl-2/Bax ratio was significantly decreased at both 24 h and 72 h after reperfusion (P<0.05). Compared with the NG group, the HG group exhibited significantly increased expression levels of Cleaved Caspase-3 and Cyt-c, and a decreased Bcl-2/Bax ratio at 24 h after I/R (P<0.05). Compared with the HG group, the HG+SE group showed a significant decrease in the Cleaved Caspase-3/Caspase-3 ratio at 24 h after I/R, and a significant reduction in both the Cleaved caspase-3/Caspase-3 ratio and Cyt-c expression levels at 72 h after I/R (P<0.05). The Bcl-2/Bax ratio was significantly elevated in the HG+SE group at both time points (P<0.05).

### 3.5 SE alleviates apoptosis-related factor expression in the penumbra of cerebral I/R rats with hyperglycemia

Immunohistochemical analysis of Fas, FasL, and PUMA factors in the penumbra of rat brains is shown in Fig 5a-f. Compared with the Sham group, the number of positive cells in the NG group, HG group, and HG+SE group significantly increased (P<0.05). At 24 h and 72 h after I/R, the number of positive cells in the HG group was significantly higher than in the NG group (P<0.05). Compared with the HG group, the HG+SE group exhibited fewer positive cells at both 24 h and 72 h (P<0.05).

**Fig 5.**
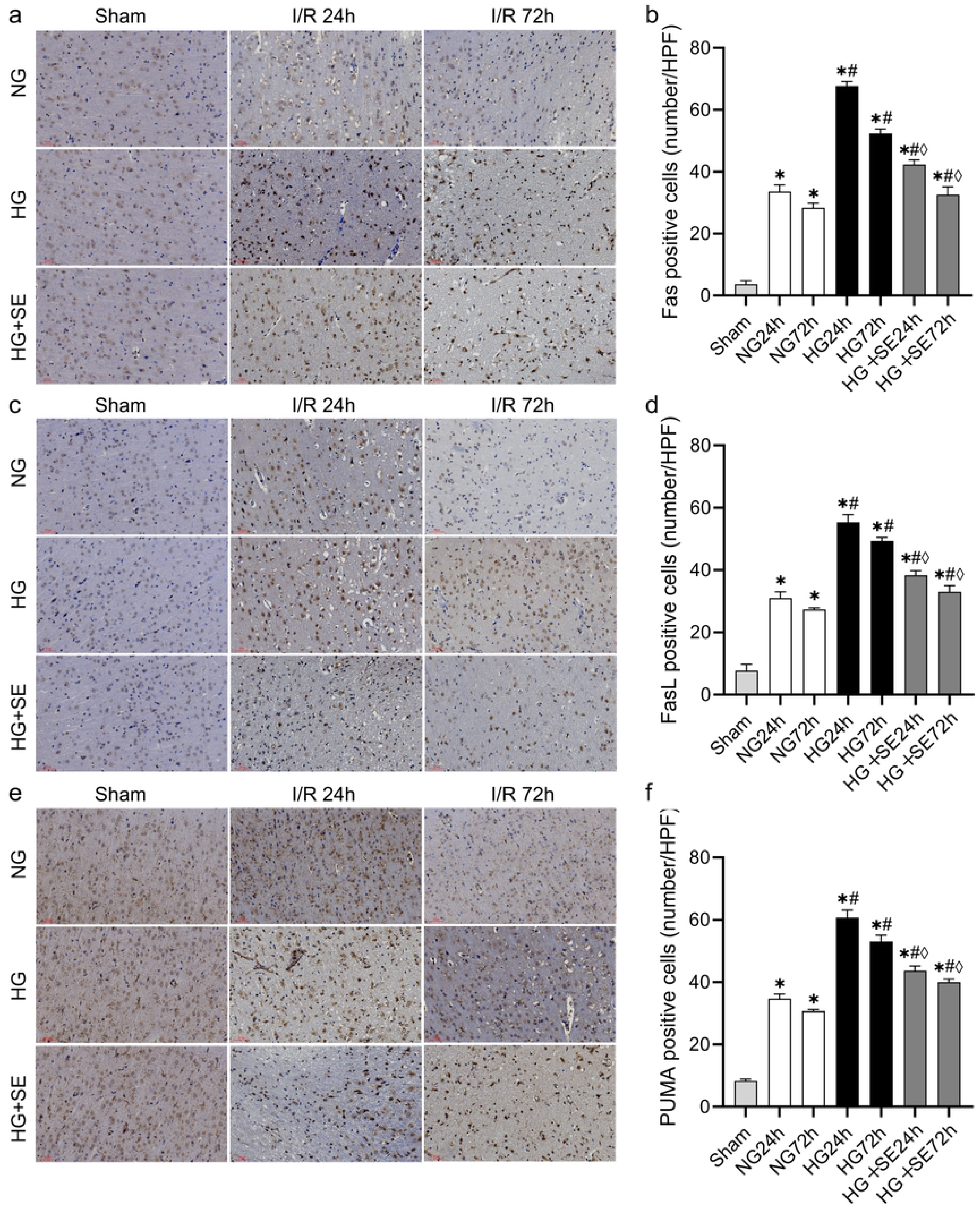
SE alleviates the expression of apoptotic factors in the penumbra of cerebral I/R rats aggravated by hyperglycemia. **a** Immunohistochemical staining of Fas-positive cells in ischemic penumbra (scale bar = 50 μm). **b** Quantitative analysis of Fas-positive cells from a (n=3). **c** Immunohistochemical staining of FasL-positive cells in ischemic penumbra (scale bar = 50 μm). **d** Quantitative analysis of FasL-positive cells from c (n=3). **e** Immunohistochemical staining of PUMA-positive cells in ischemic penumbra (scale bar = 50 μm). **f** Quantitative analysis of PUMA-positive cells from e (n=3). *P < 0.05, compared with the Sham group; ^#^P < 0.05, compared with the NG+I/R group; ^◊^P < 0.05, compared with the HG+I/R group.

### 3.6 SE slleviates high-glucose-exacerbated OD/R injury in HT22 cells

Cells were cultured under optimized conditions: OD duration (1h), glucose concentration (50 mmol/L), and SE intervention concentration (100 nmol/L). As shown in Fig 6a, the CON group displayed abundant cells with normal morphology, rounded nuclei, and intact dendritic branches. The NG+OD/R group showed reduced cell numbers, slight shrinkage, and increased intercellular spacing. The HG+OD/R group exhibited a dramatic reduction in normal cell numbers, low cell density, and further increased cell shrinkage. The HG+OD/R+SE group demonstrated improved morphological changes and increased normal cell numbers. The HG+OD/R+SE+3MA group showed further improvement in both cell numbers and morphology.

**Fig 6.**
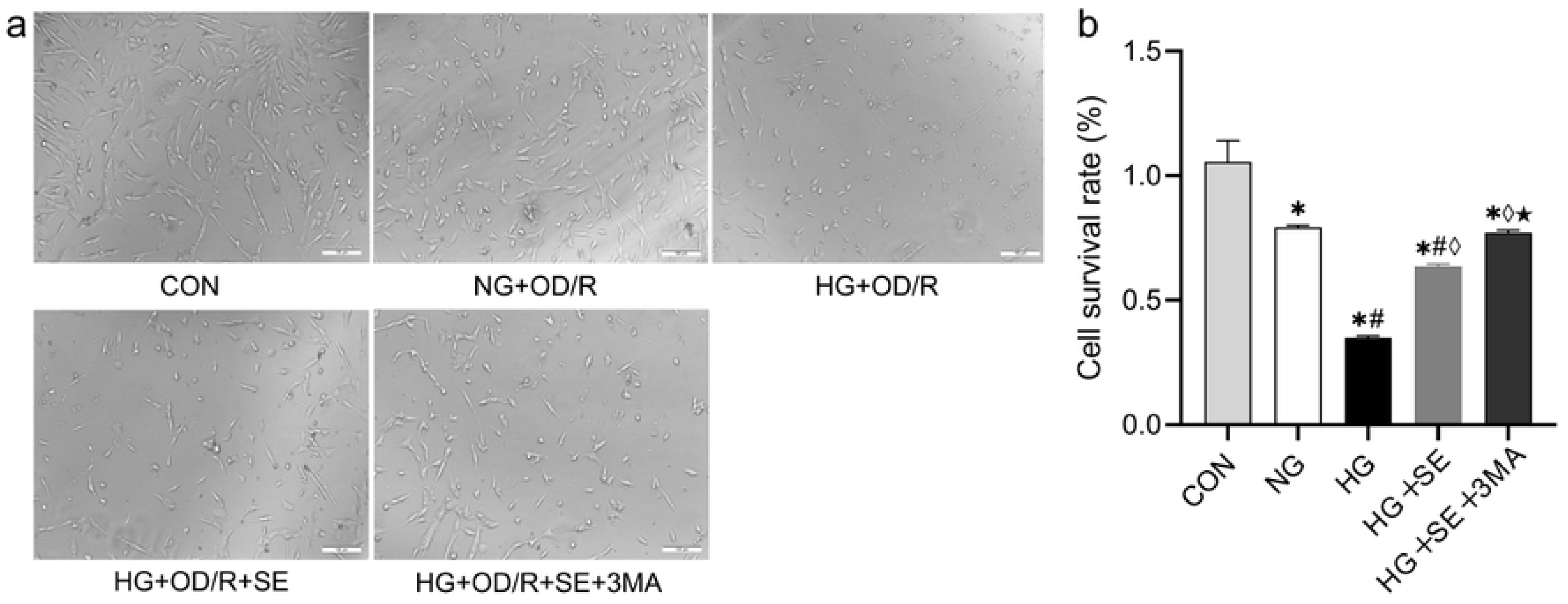
SE alleviates the damage of HT22 cells induced by high-glucose and OD/R. **a** Light microscopy imaging of HT22 cells (scale bar = 100 μm). **b** Quantitative analysis of cell viability by CCK-8 assay (n=3). *P < 0.05, compared with the CON group; ^#^P < 0.05, compared with the NG+OD/R group; ^◊^P < 0.05, compared with the HG+OD/R group; ^★^P < 0.05, compared with the HG+OD/R+SE group.

CCK-8 assay results (Fig 6b) revealed that the CON group had the highest cell viability. Compared with the CON group, the NG+OD/R group, HG+OD/R group, HG+OD/R+SE group, and HG+OD/R+SE+3MA group all showed significantly decreased cell viability (P<0.05). Compared with the NG+OD/R group, cell viability was further reduced in the HG+OD/R group and HG+OD/R+SE group (P<0.05). Compared with the HG+OD/R group, cell viability was significantly increased in the HG+OD/R+SE group and HG+OD/R+SE+3MA group (P<0.05). Additionally, a significant difference was observed between the HG+OD/R+SE+3MA group and the HG+OD/R+SE group (P<0.05).

### 3.7 SE alleviates High-glucose-aggravated HT22 cell injury via suppressing Fas/FasL pathway and apoptosis-related factors

The expression of Fas, FasL, PUMA, NF-κB, p-NF-κB and apoptosis-related factors in HT22 cells is shown in Fig 7. In Figs. 7a-f, compared with the CON group, the expression levels of Fas, FasL, PUMA and p-NF-κB in the NG+OD/R group, HG+OD/R group, and HG+OD/R+SE group were significantly increased (P<0.05). Compared with the NG+OD/R group, the expression levels of Fas, FasL and PUMA in the HG+OD/R group were significantly elevated (P < 0.05). Compared with the HG+OD/R group, the expression levels of Fas, FasL and PUMA in the HG+OD/R+SE group and HG+OD/R+SE+3MA were significantly reduced (P<0.05). The HG+OD/R+SE+3MA group exhibited more significant decreases in the p-NF-κB/NF-κB ratio and PUMA expression compared with the HG+OD/R group (P<0.05).

**Fig 7.**
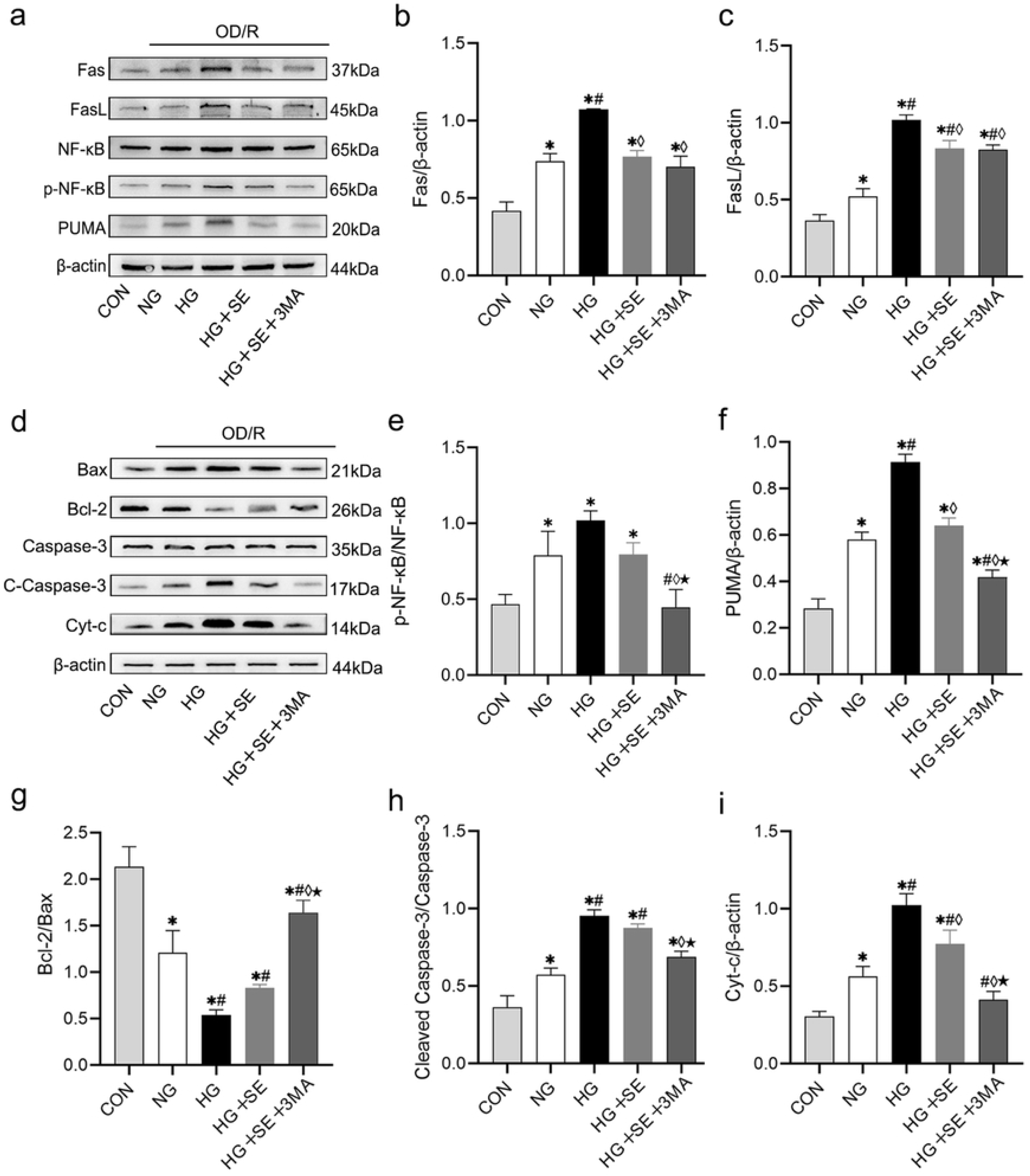
SE alleviates the damage of HT22 cells induced by high-glucose OD/R by reducing the expression of factors in the Fas/FasL pathway and apoptotic factors. **a** Representative Western Blot results of Fas/FasL pathway proteins in HT22 cells. **b** Quantitative analysis of Fas expression (relative to β-actin) from a (n=3). **c** Quantitative analysis of FasL expression (relative to β-actin) from a (n=3). **e** Quantitative analysis of NF-κB activation level (p-NF-κB/NF-κB ratio) from a (n=3). **f** Quantitative analysis of PUMA expression (relative to β-actin) from a (n=3). **d** Representative Western Blot results of apoptosis related proteins in HT22 cells. **g** Quantitative analysis of Bcl-2/Bax expression from d (n=3). **h** Quantitative analysis of Cleaved Caspase-3/Caspase-3 expression from d (n=3). **i** Quantitative analysis of Cyt-c (relative to β-actin) from d (n=3). *P < 0.05, compared with the CON group; ^#^P < 0.05, compared with the NG+OD/R group; ^◊^P < 0.05, compared with the HG+OD/R group; ^★^P < 0.05, compared with the HG+OD/R+SE group.

Figs. 7d/g/h/i show apoptosis-related factors (Bax, Bcl-2, Cleaved Caspase-3 and Cyt-c) in HT22 cells. In Figs. 7g/h/i, compared with the CON group, the expression levels of Cleaved Caspase-3 and Cyt-c in the NG+OD/R, HG+OD/R and HG+OD/R+SE groups were significantly increased (P<0.05), while the CON group showed the highest Bcl-2/Bax ratio (P<0.05). Compared with the NG+OD/R group, both the HG+OD/R and HG+OD/R+SE groups displayed significantly decreased Bcl-2/Bax ratios (P<0.05), along with significantly elevated Cleaved Caspase-3/Caspase-3 ratios and Cyt-c expression (P<0.05). Compared with the HG+OD/R+SE group, the HG+OD/R+SE+3MA group showed a significantly increased Bcl-2/Bax ratio (P<0.05), and reduced Cleaved Caspase-3/Caspase-3 ratio and Cyt-c expression (P<0.05).

### 3.8 Apoptosis-related factor expression in PUMA-knockdown HT22 cells

As shown in Figs. 8a-b, the PUMA knockdown group (KD) exhibited significantly decreased PUMA expression compared with wild-type cells (WT) (P<0.05). Figs. 8c-i demonstrate that in PUMA-WT cells: Compared with the CON group, the expression levels of Bax, Cleaved Caspase-3, Cyt-c and PUMA were significantly increased in both the HG+OD/R and HG+OD/R+SE groups (P<0.05). The HG+OD/R+SE group showed significantly reduced expression of these factors compared with the HG+OD/R group (P<0.05). In PUMA-KD cells: Both the HG+OD/R and HG+OD/R+SE groups showed decreasing trends in Bax, Cleaved Caspase-3, Cyt-c and PUMA expression compared with WT cells. Specifically, the HG+OD/R+SE group in PUMA-KD cells exhibited significantly lower expression of Bax, Cleaved Caspase-3, Cyt-c and PUMA compared with WT cells (P<0.05).

**Fig 8.**
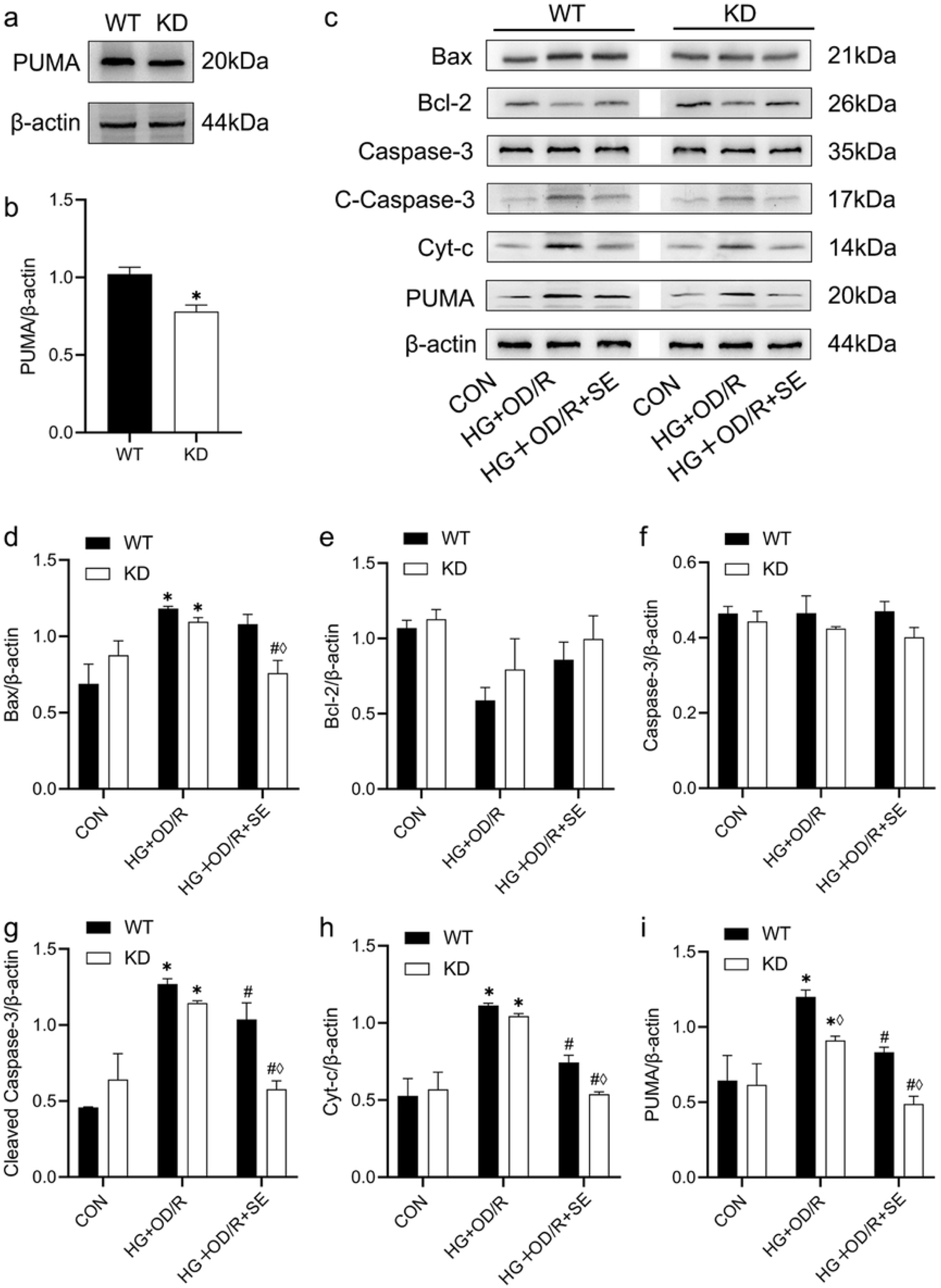
Expression of apoptosis factors after PUMA knockdown in HT22 cells. **a** Representative Western Blot validation of PUMA knockdown in HT22 cells. **b** Quantitative analysis of PUMA knockdown efficiency (relative to β-actin) from a (n=3). **c** Representative Western Blot results of apoptosis-related proteins post-PUMA knockdown. **d** Quantitative analysis of Bax expression (relative to β-actin) from c (n=3). **e** Quantitative analysis of Bcl-2 expression (relative to β-actin) from c (n=3). **f** Quantitative analysis of Caspase-3 expression (relative to β-actin) from c (n=3). **g** Quantitative analysis of Cleaved Caspase-3 expression (relative to β-actin) from c (n=3). **h** Quantitative analysis of Cyt-c expression (relative to β-actin) from c (n=3). **i** Quantitative analysis of residual PUMA expression (relative to β-actin) from c (n=3). *P < 0.05, compared with the CON group; ^#^P < 0.05, compared with the HG+OD/R group; ^◊^P < 0.05, compared with WT-HT22 cells under the same intervention conditions.

## 4. Discussion

Clinical data show that upon admission, two-thirds of patients with ischemic stroke have blood glucose levels higher than 6.0 mmol/L [39]. The exacerbation of ischemic stroke injury caused by hyperglycemia involves many mechanisms, including lactate accumulation, inflammatory response, and production of reactive oxygen species (ROS) **Error! Reference source not found. Error! Reference source not found.**. Macrovascular diseases and microvascular reactive injuries caused by diabetes are also important mechanisms leading to brain injury [40] [43]. In this study, the neurological deficit score in the HG group was significantly higher than that in the NG group, accompanied by aggravated neuronal edema and an increased proportion of nuclear pyknosis, suggesting that hyperglycemia can significantly amplify ischemic injury. This finding is consistent with previous research conclusions, that is, the adverse cerebrovascular remodeling and abnormal perfusion caused by hyperglycemia will exacerbate hypoxia in brain tissue after stroke [44][45].

SE, as an essential trace element [46], shows the potential for multi-target intervention. Studies have shown that SE can reduce oxidative stress, apoptosis, and inflammation in damaged brain tissue by mediating multiple signaling pathways [47]. Due to its antioxidant stress activity and regulation of glucose transport and glucose metabolism, SE may have potential therapeutic applications in the treatment of diseases such as diabetes and brain injury **Error! Reference source not found.**. This study also verified the intervention effect of SE. In the HG group, food intake, water intake, and urine output increased significantly, and the HG+SE group could partially improve the symptoms of polydipsia and polyuria, HG increased the infarct volume, neurological deficit, and neuronal apoptosis in the penumbra area of rats with cerebral ischemia, while the brain injury in the HG+SE group was milder. The in-vitro experimental results were generally consistent with the in-vivo research results: high-glucose OD caused morphological damage to HT22 cells, decreased cell viability, and increased the expression of apoptotic factors, and SE could improve these manifestations. The above results are consistent with the previously reported mechanism of SE improving insulin sensitivity [49]**Error! Reference source not found.**.

Another study revealed that SE can reduce apoptosis of spinal ganglion and hippocampal neurons in diabetic rats [50]. Based on the above research reports, this study focused on cell apoptosis. Cell apoptosis is a highly regulated cell death mechanism, and its core execution system depends on the cascade activation of the Caspase family of proteases and the precise regulation of mitochondrial membrane permeability by the Bcl-2 protein family. When cells are stimulated by endogenous or exogenous factors such as DNA damage, hypoxia, or abnormal expression of oncoproteins (such as Myc, Ras), apoptotic signals are transmitted through two main pathways: the exogenous death receptor pathway is triggered by the binding of Fas/TNF receptors on the cell membrane to their ligands, recruits the adaptor protein FADD, activates Caspase-8, and then directly cleaves the downstream executioner Caspases-3/7 or generates the active fragment tBid by cleaving Bid, which translocates to the mitochondria to activate the endogenous pathway; the endogenous mitochondrial pathway forms pores on the outer mitochondrial membrane through the oligomerization of the pro-apoptotic protein Bax, leading to the release of Cyt-c and its binding to Apaf-1 in the cytoplasm to form an apoptosome, activating Caspase-9, and then initiating the cell disintegration program mediated by Caspases-3/7 [52]**Error! Reference source not found.**. In this process, the anti-apoptotic protein Bcl-2 maintains mitochondrial homeostasis and blocks the transmission of apoptotic signals by competitively binding to Bax or inhibiting mitochondrial membrane permeability transition (MPTP). The final stage of apoptosis is dominated by activated Caspases-3/7, which triggers chromatin condensation and nuclear fragmentation by cleaving key substrates such as nuclear lamin and PARP, and finally forms membrane-wrapped apoptotic bodies. The phosphatidylserine (PS) exposed on their surface is recognized and cleared by macrophages as a "phagocytic signal" to avoid an inflammatory response **Error! Reference source not found.**. This precisely regulated network not only involves the dynamic balance of the Caspase and Bcl-2 families but is also cross-regulated by signaling pathways such as NF-κB and p53, ensuring that the organism achieves overall homeostasis by selectively eliminating abnormal cells during development, immunity, and pathological states.

The NF-κB signaling pathway, as the core hub of apoptosis regulation, is not only involved in physiological processes such as cell growth and differentiation but also plays a key role in cell death through the Fas/FasL/NF-κB/PUMA cascade reaction **Error! Reference source not found.**. In pathological states such as cerebral ischemia, the death receptor Fas is activated due to the up-regulation of its ligand FasL. This process starts within 12 h after ischemia and peaks at 24-48 h, which is highly synchronized with the process of neuronal apoptosis [56]**Error! Reference source not found.**. The activated Fas receptor forms a death-inducing signaling complex (DISC) by recruiting FADD and Caspase-8, directly triggering the apoptosis execution program; at the same time, the Fas signal can also regulate downstream target genes by activating NF-κB p65. After NF-κB dissociates from IκB, it quickly enters the nucleus and induces the transcription of the pro-apoptotic factor PUMA (p53 up-regulated modulator of apoptosis) [59][61]. As an important part of the apoptosis process, PUMA promotes the oligomerization of pro-apoptotic proteins Bax/Bak on the mitochondrial membrane by antagonizing anti-apoptotic proteins Bcl-2, Mcl-1, and Bcl-xL, leading to the release of Cyt-c and activation of Caspase-9, and then driving mitochondrial-dependent apoptosis **Error! Reference source not found.Error! Reference source not found.**. This study revealed that compared with the NG group, the expression levels of Fas, FasL, and PUMA and the phosphorylation of NF-κB in the brain tissue of rats in the HG group were significantly increased, indicating that hyperglycemia exacerbates cerebral I/R injury by activating the Fas/FasL/NF-κB/PUMA signaling pathway. When the PUMA gene was knocked down, the expression of pro-apoptotic factors such as Bax, Cleaved Caspase-3, and Cyt-c was inhibited, confirming that PUMA is the core hub regulating mitochondrial-dependent apoptosis. This result is consistent with previous studies on the molecular mechanisms: Fas/FasL regulates PUMA transcription through the NF-κB signaling axis [59][61], and PUMA promotes mitochondrial apoptosis by antagonizing Bcl-2 family proteins (such as Bcl-2 and Mcl-1) **Error! Reference source not found.Error! Reference source not found.**. The research data further reveals that SE intervention can inhibit the expression of apoptotic molecules, reduce the volume of cerebral infarction and the structural damage of neurons. By using Fas and FasL inhibitors and knocking down the expression of PUMA, it is shown that SE can regulate the PUMA apoptotic pathway through Fas/FasL, inhibit the expression of pro-apoptotic factors such as Bax, Cleaved Caspase-3 and Cyt-c, and alleviate cerebral I/R injury, providing a new strategy for the intervention and treatment of hyperglycemia complicated with cerebral ischemic injury.

In conclusion, this study confirms that SE can alleviate cerebral I/R injury aggravated by hyperglycemia by regulating apoptosis mediated by the Fas/FasL/NF-κB/PUMA signaling pathway. Our findings on the relationship between SE, the Fas/FasL/NF-κB/PUMA signaling pathway and neuroprotection will contribute to the exploration of new therapeutic targets and drugs for cerebral I/R injury aggravated by hyperglycemia. To benefit diabetic patients and patients with ischemic stroke, it is urgent to conduct further clinical trials to verify our observations.

## Author Contributions

**Conceptualization**: Feng Ding, Xida Yin.

**Methodology**: Lan Yang.

**Investigation**: Feng Ding, Bowen Zheng.

**Resources**: Lan Yang, Jingwen Zhang.

**Writing–original draft**: Shuai Zhao.

**Writing–review & editing**: Feng Ding, Xida Yin.

**Visualization**:Yue Chang.

**Funding acquisition:**Lan Yang, Jingwen Zhang.

## References

[1] Katan M, Luft A. Global Burden of Stroke. Semin Neurol. 2018 Apr;38(2):208–211. doi: 10.1055/s-0038-1649503. Epub 2018 May 23. PMID: 29791947.

[2] Paul S, Candelario-Jalil E. Emerging neuroprotective strategies for the treatment of ischemic stroke: An overview of clinical and preclinical studies. Exp Neurol. 2021 Jan;335:113518. doi: 10.1016/j.expneurol.2020.113518. Epub 2020 Nov 2. PMID: 33144066; PMCID: PMC7869696.

[3] Akinyemi RO, Ovbiagele B, Adeniji OA, Sarfo FS, Abd-Allah F, Adoukonou T, et al. Stroke in Africa: profile, progress, prospects and priorities. Nat Rev Neurol. 2021 Oct;17(10):634–656. doi: 10.1038/s41582-021-00542-4. Epub 2021 Sep 15. PMID: 34526674; PMCID: PMC8441961.

[4] Kim J, Olaiya MT, De Silva DA, Norrving B, Bosch J, De Sousa DA, et al. Global stroke statistics 2023: Availability of reperfusion services around the world. Int J Stroke. 2024 Mar;19(3):253-270. doi: 10.1177/17474930231210448. Epub 2024 Jan 1. PMID: 37853529; PMCID: PMC10903148.

[5] Lin W, Zhao XY, Cheng JW, Li LT, Jiang Q, Zhang YX, et al. Signaling pathways in brain ischemia: Mechanisms and therapeutic implications. Pharmacol Ther. 2023 Nov;251:108541. doi: 10.1016/j.pharmthera.2023.108541. Epub 2023 Oct 1. PMID: 37783348.

[6] Li X, Ma N, Xu J, Zhang Y, Yang P, Su X, et al.Targeting Ferroptosis: Pathological Mechanism and Treatment of Ischemia-Reperfusion Injury. Oxid Med Cell Longev. 2021 Oct 28;2021:1587922. doi: 10.1155/2021/1587922. PMID: 34745412; PMCID: PMC8568519.

[7] Pan J, Konstas AA, Bateman B, Ortolano GA, Pile-Spellman J. Reperfusion injury following cerebral ischemia: pathophysiology, MR imaging, and potential therapies. Neuroradiology. 2007 Feb;49(2):93–102. doi: 10.1007/s00234-006-0183-z. Epub 2006 Dec 20. PMID: 17177065; PMCID: PMC1786189.

[8] Anderson CS. Progress-Defining Risk Factors for Stroke Prevention. Cerebrovasc Dis. 2021;50(6):615–616. doi: 10.1159/000516996. Epub 2021 May 27. PMID: 34044390.

[9] Yang J, Liu Z. Mechanistic Pathogenesis of Endothelial Dysfunction in Diabetic Nephropathy and Retinopathy. Front Endocrinol (Lausanne). 2022 May 25;13:816400. doi: 10.3389/fendo.2022.816400. PMID: 35692405; PMCID: PMC9174994.

[10] Reske-Nielsen E, Lundbæk K, Rafaelsen OJ. Pathological changes in the central and peripheral nervous system of young long-term diabetics : I. Diabetic encephalopathy. Diabetologia. 1966 Apr;1(3-4):233–41. doi: 10.1007/BF01257917. PMID: 24173307.

[11] Grunnet ML. Cerebrovascular disease: diabetes and cerebral atherosclerosis. Neurology. 1963 Jun;13:486–91. doi: 10.1212/wnl.13.6.486. PMID: 13950949.

[12] Tu WJ, Wang LD; Special Writing Group of China Stroke Surveillance Report. China stroke surveillance report 2021. Mil Med Res. 2023 Jul 19;10(1):33. doi: 10.1186/s40779-023-00463-x. PMID: 37468952; PMCID: PMC10355019.

[13] Neves D, Salazar IL, Almeida RD, Silva RM. Molecular mechanisms of ischemia and glutamate excitotoxicity. Life Sci. 2023 Sep 1;328:121814. doi: 10.1016/j.lfs.2023.121814. Epub 2023 May 24. PMID: 37236602.

[14] Moawad MHED, Salem T, Alaaeldin A, Elaraby Y, Awad PD, Khalifa AA, et al. Safety and efficacy of intravenous thrombolysis: a systematic review and meta-analysis of 93,057 minor stroke patients. BMC Neurol. 2025 Jan 22;25(1):33. doi: 10.1186/s12883-024-04000-8. PMID: 39844066; PMCID: PMC11752810.

[15] Paskiewicz A, Niu J, Chang C. Autoimmune lymphoproliferative syndrome: A disorder of immune dysregulation. Autoimmun Rev. 2023 Nov;22(11):103442. doi: 10.1016/j.autrev.2023.103442. Epub 2023 Sep 6. PMID: 37683818.

[16] Chelluboina B, Klopfenstein JD, Gujrati M, Rao JS, Veeravalli KK. Temporal regulation of apoptotic and anti-apoptotic molecules after middle cerebral artery occlusion followed by reperfusion. Mol Neurobiol. 2014 Feb;49(1):50–65. doi: 10.1007/s12035-013-8486-7. Epub 2013 Jun 28. PMID: 23813097; PMCID: PMC3918127.

[17] Muhammad IF, Borné Y, Melander O, Orho-Melander M, Nilsson J, Söderholm M, et al. FADD (Fas-Associated Protein With Death Domain), Caspase-3, and Caspase-8 and Incidence of Ischemic Stroke. Stroke. 2018 Sep;49(9):2224-2226. doi: 10.1161/STROKEAHA.118.022063. PMID: 30354994.

[18] Alharbi KS, Fuloria NK, Fuloria S, Rahman SB, Al-Malki WH, Javed Shaikh MA, et al. Nuclear factor-kappa B and its role in inflammatory lung disease. Chem Biol Interact. 2021 Aug 25;345:109568. doi: 10.1016/j.cbi.2021.109568. Epub 2021 Jun 25. PMID: 34181887.

[19] Yang S, Zhang X, Qu H, Qu B, Yin X, Zhao H. Retraction Note: Cabozantinib induces PUMA-dependent apoptosis in colon cancer cells via AKT/GSK-3β/NF-κB signaling pathway. Cancer Gene Ther. 2022 Nov;29(11):1806. doi: 10.1038/s41417-022-00545-3. PMID: 36241704; PMCID: PMC9663299.

[20] Singh R, Yu S, Osman M, Inde Z, Fraser C, Cleveland AH, et al. Radiotherapy-Induced Neurocognitive Impairment Is Driven by Heightened Apoptotic Priming in Early Life and Prevented by Blocking BAX. Cancer Res. 2023 Oct 13;83(20):3442–3461. doi: 10.1158/0008-5472.CAN-22-1337. PMID: 37470810; PMCID: PMC10570680.

[21] Su Y, Lu S, Hou C, Ren K, Wang M, Liu X, et al. Mitigation of liver fibrosis via hepatic stellate cells mitochondrial apoptosis induced by metformin. Int Immunopharmacol. 2022 Jul;108:108683. doi: 10.1016/j.intimp.2022.108683. Epub 2022 Mar 25. PMID: 35344814.

[22] Tan S, Liu X, Chen L, Wu X, Tao L, Pan X, et al. Fas/FasL mediates NF-κBp65/PUMA-modulated hepatocytes apoptosis via autophagy to drive liver fibrosis. Cell Death Dis. 2021 May 12;12(5):474. doi: 10.1038/s41419-021-03749-x. PMID: 33980818; PMCID: PMC8115181.

[23] Kumar A, Prasad KS. Role of nano-selenium in health and environment. J Biotechnol. 2021 Jan 10;325:152–163. doi: 10.1016/j.jbiotec.2020.11.004. Epub 2020 Nov 4. PMID: 33157197.

[24] Yang H, Wang Z, Li L, Wang X, Wei X, Gou S, et al. Mannose coated selenium nanoparticles normalize intestinal homeostasis in mice and mitigate colitis by inhibiting NF-κB activation and enhancing glutathione peroxidase expression. J Nanobiotechnology. 2024 Oct 10;22(1):613. doi: 10.1186/s12951-024-02861-2. PMID: 39385176; PMCID: PMC11465824.

[25] Pitts MW, Hoffmann PR, Schomburg L. Editorial: Selenium and Selenoproteins in Brain Development, Function, and Disease. Front Neurosci. 2022 Jan 13;15:821140. doi: 10.3389/fnins.2021.821140. PMID: 35095409; PMCID: PMC8792733.

[26] Zhang F, Li X, Wei Y. Selenium and Selenoproteins in Health. Biomolecules. 2023 May 8;13(5):799. doi: 10.3390/biom13050799. PMID: 37238669; PMCID: PMC10216560.

[27] Chen YX, Zuliyaer T, Liu B, Guo S, Yang DG, Gao F, et al. Sodium selenite promotes neurological function recovery after spinal cord injury by inhibiting ferroptosis. Neural Regen Res. 2022 Dec;17(12):2702–2709. doi: 10.4103/1673-5374.339491. PMID: 35662217; PMCID: PMC9165358.

[28] Yang L, Ma YM, Shen XL, Fan YC, Zhang JZ, Li PA, et al. The Involvement of Mitochondrial Biogenesis in Selenium Reduced Hyperglycemia-Aggravated Cerebral Ischemia Injury. Neurochem Res. 2020 Aug;45(8):1888–1901. doi: 10.1007/s11064-020-03055-6. Epub 2020 May 23. PMID: 32447509.

[29] Raza A, Johnson H, Singh A, Sharma AK. Impact of selenium nanoparticles in the regulation of inflammation. Arch Biochem Biophys. 2022 Dec 15;732:109466. doi: 10.1016/j.abb.2022.109466. Epub 2022 Nov 17. PMID: 36403759.

[30] Stachowicz K. Regulation of COX-2 expression by selected trace elements and heavy metals: Health implications, and changes in neuronal plasticity. A review. J Trace Elem Med Biol. 2023 Sep;79:127226. doi: 10.1016/j.jtemb.2023.127226. Epub 2023 May 25. PMID: 37257334.

[31] Li G, Cheng Y, Ding S, Zheng Q, Kuang L, Zhang Y, Zhou Y. Identification of key genes associated with oxidative stress in ischemic stroke via bioinformatics integrated analysis. BMC Neurosci. 2025 Jan 13;26(1):3. doi: 10.1186/s12868-024-00921-9. PMID: 39806309; PMCID: PMC11727628.

[32] Albert-Garay JS, Riesgo-Escovar JR, Salceda R. High glucose concentrations induce oxidative stress by inhibiting Nrf2 expression in rat Müller retinal cells in vitro. Sci Rep. 2022 Jan 24;12(1):1261. doi: 10.1038/s41598-022-05284-x. PMID: 35075205; PMCID: PMC8975969.

[33] Sohail A, Murtaza Hasnain M, Ehsan Ul Haq M, Nasir I, Sufyan R, Khan M, et al. Oxidative Stress and Antioxidant Interventions in Type 2 Diabetes [Internet]. Biochemistry. IntechOpen; 2024. Available from: 10.5772/intechopen.1006081

[34] Abu Khadra KM, Bataineh MI, Khalil A, Saleh J. Oxidative stress and type 2 diabetes: the development and the pathogenesis, Jordanian cross-sectional study. Eur J Med Res. 2024 Jul 17;29(1):370. doi: 10.1186/s40001-024-01906-4. PMID: 39014510; PMCID: PMC11253486.

[35] Mehta SL, Kumari S, Mendelev N, Li PA. Selenium preserves mitochondrial function, stimulates mitochondrial biogenesis, and reduces infarct volume after focal cerebral ischemia. BMC Neurosci. 2012 Jul 9;13:79. doi: 10.1186/1471-2202-13-79. PMID: 22776356; PMCID: PMC3411431.

[36] Shalihat A, Hasanah AN, Mutakin, Lesmana R, Budiman A, Gozali D. The role of selenium in cell survival and its correlation with protective effects against cardiovascular disease: A literature review. Biomed Pharmacother. 2021 Feb;134:111125. doi: 10.1016/j.biopha.2020.111125. Epub 2020 Dec 16. PMID: 33341057.

[37] Huang Y, Ni Y, Yu L, Shu L, Zhu Q, He X. Dietary total antioxidant capacity and risk of stroke: a systematic review and dose-response meta-analysis of observational studies. Front Nutr. 2024 Sep 19;11:1451386. doi: 10.3389/fnut.2024.1451386. PMID: 39364151; PMCID: PMC11448356.

[38] Pawluk H, Tafelska-Kaczmarek A, Sopońska M, Porzych M, Modrzejewska M, Pawluk M, et al. The Influence of Oxidative Stress Markers in Patients with Ischemic Stroke. Biomolecules. 2024 Sep 6;14(9):1130. doi: 10.3390/biom14091130. PMID: 39334896; PMCID: PMC11430825.

[39] Feske SK. Ischemic Stroke. Am J Med. 2021 Dec;134(12):1457-1464. doi: 10.1016/j.amjmed.2021.07.027. Epub 2021 Aug 27. PMID: 34454905.

[40] Nogueira RG, Jadhav AP, Haussen DC, Bonafe A, Budzik RF, Bhuva P, et al. Thrombectomy 6 to 24 Hours after Stroke with a Mismatch between Deficit and Infarct. N Engl J Med. 2018 Jan 4;378(1):11–21. doi: 10.1056/NEJMoa1706442. Epub 2017 Nov 11. PMID: 29129157.

[41] Rehni AK, Cho S, Dave KR. Ischemic brain injury in diabetes and endoplasmic reticulum stress. Neurochem Int. 2022 Jan;152:105219. doi: 10.1016/j.neuint.2021.105219. Epub 2021 Nov 1. PMID: 34736936; PMCID: PMC8918032.

[42] González P, Lozano P, Ros G, Solano F. Hyperglycemia and Oxidative Stress: An Integral, Updated and Critical Overview of Their Metabolic Interconnections. Int J Mol Sci. 2023 May 27;24(11):9352. doi: 10.3390/ijms24119352. PMID: 37298303; PMCID: PMC10253853.

[43] Hu X, De Silva TM, Chen J, Faraci FM. Cerebral Vascular Disease and Neurovascular Injury in Ischemic Stroke. Circ Res. 2017 Feb 3;120(3):449–471. doi: 10.1161/CIRCRESAHA.116.308427. PMID: 28154097; PMCID: PMC5313039.

[44] Arcambal A, Taïlé J, Couret D, Planesse C, Veeren B, Diotel N, et al. Protective Effects of Antioxidant Polyphenols against Hyperglycemia-Mediated Alterations in Cerebral Endothelial Cells and a Mouse Stroke Model. Mol Nutr Food Res. 2020 Jul;64(13):e1900779. doi: 10.1002/mnfr.201900779. Epub 2020 Jun 16. PMID: 32447828.

[45] Jiang Y, Han J, Li Y, Wu Y, Liu N, Shi SX, et al. Delayed rFGF21 Administration Improves Cerebrovascular Remodeling and White Matter Repair After Focal Stroke in Diabetic Mice. Transl Stroke Res. 2022 Apr;13(2):311–325. doi: 10.1007/s12975-021-00941-1. Epub 2021 Sep 15. PMID: 34523038.

[46] Poittevin M, Bonnin P, Pimpie C, Rivière L, Sebrié C, Dohan A, et al. Diabetic microangiopathy: impact of impaired cerebral vasoreactivity and delayed angiogenesis after permanent middle cerebral artery occlusion on stroke damage and cerebral repair in mice. Diabetes. 2015 Mar;64(3):999–1010. doi: 10.2337/db14-0759. Epub 2014 Oct 6. PMID: 25288671.

[47] Xiang H, Tan Q, Zhang Y, Wu Y, Xu Y, Hong Y, et al. Sodium selenite attenuates inflammatory response and oxidative stress injury by regulating the Nrf2/ARE pathway in contrast-induced acute kidney injury in rats. BMC Nephrol. 2024 Jul 15;25(1):226. doi: 10.1186/s12882-024-03657-0. PMID: 39009991; PMCID: PMC11247789.

[48] Li LX, Chu JH, Chen XW, Gao PC, Wang ZY, Liu C, et al. Selenium ameliorates mercuric chloride-induced brain damage through activating BDNF/TrKB/PI3K/AKT and inhibiting NF-κB signaling pathways. J Inorg Biochem. 2022 Apr;229:111716. doi: 10.1016/j.jinorgbio.2022.111716. Epub 2022 Jan 5. PMID: 35065321.

[49] Wang X, Sui H, Su Y, Zhao S. Protective effects of sodium selenite on insulin secretion and diabetic retinopathy in rats with type 1 diabetes mellitus. Pak J Pharm Sci. 2021 Sep;34(5):1729–1735. PMID: 34803009.

[50] Han J, Wu Y, Wang Z, Han J, Luo G, Huo K. Early venous filling is associated with unfavorable outcomes in acute ischemic stroke with large vessel occlusion after mechanical thrombectomy: a real-world analysis. BMC Neurol. 2025 Mar 6;25(1):92. doi: 10.1186/s12883-025-04111-w. PMID: 40050750; PMCID: PMC11883998.

[51] Xu X, Qi P, Zhang Y, Sun H, Yan Y, Sun W, Liu S. Effect of Selenium Treatment on Central Insulin Sensitivity: A Proteomic Analysis in β-Amyloid Precursor Protein/Presenilin-1 Transgenic Mice. Front Mol Neurosci. 2022 Jul 7;15:931788. doi: 10.3389/fnmol.2022.931788. PMID: 35875664; PMCID: PMC9302600.

[52] Aliakbari K, Saidie P. High intensity interval training and selenium nanoparticles protect hippocampal neurons and enhance cognitive function in diabetic rats. Sci Rep. 2025 Jul 1;15(1):21531. doi: 10.1038/s41598-025-07441-4. PMID: 40593054; PMCID: PMC12216540.

[53] Kerr JF. History of the events leading to the formulation of the apoptosis concept. Toxicology. 2002 Dec 27;181–182:471-4. doi: 10.1016/s0300-483x(02)00457-2. PMID: 12505355.

[54] Cohen GM. Caspases: the executioners of apoptosis. Biochem J. 1997 Aug 15;326 ( Pt 1)(Pt 1):1-16. doi: 10.1042/bj3260001. PMID: 9337844; PMCID: PMC1218630.

[55] Levine AJ. p53, the cellular gatekeeper for growth and division. Cell. 1997 Feb 7;88(3):323–31. doi: 10.1016/s0092-8674(00)81871-1. PMID: 9039259.

[56] Tan S, Liu X, Chen L, Wu X, Tao L, Pan X, et al. Fas/FasL mediates NF-κBp65/PUMA-modulated hepatocytes apoptosis via autophagy to drive liver fibrosis. Cell Death Dis. 2021 May 12;12(5):474. doi: 10.1038/s41419-021-03749-x. PMID: 33980818; PMCID: PMC8115181.

[57] Igney FH, Krammer PH. Death and anti-death: tumour resistance to apoptosis. Nat Rev Cancer 2002 Apr;2(4):277–88. doi: 10.1038/nrc776. PMID: 12001989.

[58] Martin-Villalba A, Herr I, Jeremias I, Hahne M, Brandt R, Vogel J, et al. CD95 ligand (Fas-L/APO-1L) and tumor necrosis factor-related apoptosis-inducing ligand mediate ischemia-induced apoptosis in neurons. J Neurosci. 1999 May 15;19(10):3809–17. doi: 10.1523/JNEUROSCI.19-10-03809.1999. PMID: 10234013; PMCID: PMC6782733.

[59] Rosenbaum DM, Gupta G, D’Amore J, Singh M, Weidenheim K, Zhang H, Kessler JA. Fas (CD95/APO-1) plays a role in the pathophysiology of focal cerebral ischemia. J Neurosci Res. 2000 Sep 15;61(6):686–92. doi: 10.1002/1097-4547(20000915)61:6<686::AID-JNR12>3.0.CO;2-7. PMID: 10972965.

[60] Zinatizadeh MR, Schock B, Chalbatani GM, Zarandi PK, Jalali SA, Miri SR. The Nuclear Factor Kappa B (NF-kB) signaling in cancer development and immune diseases. Genes Dis. 2020 Jul 18;8(3):287–297. doi: 10.1016/j.gendis.2020.06.005. PMID: 33997176; PMCID: PMC8093649.

[61] Xue Q, Kang R, Klionsky DJ, Tang D, Liu J, Chen X. Copper metabolism in cell death and autophagy. Autophagy. 2023 Aug;19(8):2175–2195. doi: 10.1080/15548627.2023.2200554. Epub 2023 Apr 16. PMID: 37055935; PMCID: PMC10351475.

[62] Sun L, Huang Y, Liu Y, Zhao Y, He X, Zhang L, et al. Ipatasertib, a novel Akt inhibitor, induces transcription factor FoxO3a and NF-κB directly regulates PUMA-dependent apoptosis. Cell Death Dis. 2018 Sep 5;9(9):911. doi: 10.1038/s41419-018-0943-9. PMID: 30185800; PMCID: PMC6125489.

[63] Kurschat C, Metz A, Kirschnek S, Häcker G. Importance of Bcl-2-family proteins in murine hematopoietic progenitor and early B cells. Cell Death Dis. 2021 Aug 11;12(8):784. doi: 10.1038/s41419-021-04079-8. PMID: 34381022; PMCID: PMC8358012.

[64] Tan S, Li L, Chen T, Chen X, Tao L, Lin X, et al. β-Arrestin-1 protects against endoplasmic reticulum stress/p53-upregulated modulator of apoptosis-mediated apoptosis via repressing p-p65/inducible nitric oxide synthase in portal hypertensive gastropathy. Free Radic Biol Med. 2015 Oct;87:69–83. doi: 10.1016/j.freeradbiomed.2015.06.004. Epub 2015 Jun 25. PMID: 26119788.

[65] Thomalla D, Beckmann L, Grimm C, Oliverio M, Meder L, Herling CD, et al. Deregulation and epigenetic modification of BCL2-family genes cause resistance to venetoclax in hematologic malignancies. Blood. 2022 Nov 17;140(20):2113–2126. doi: 10.1182/blood.2021014304. PMID: 35704690; PMCID: PMC10653032.

